# Optimized antibody humanization by intra and inter VH-VL binding energy sorting

**DOI:** 10.1101/2021.01.06.425645

**Authors:** Qiang Zhong

**Author notes:** Correspondance address to Qiang Zhong.

## Abstract

Humanization of non-human derived antibody is necessary to reduce the immunogenicity of antibody drug. In recent years, computer aided antibody humanization became an efficient and rapid routine process. We report here a new computational humanization pipeline which includes CDR grafting onto the crystal structures of homologous antibody and CDR grafted humanized antibody structures. Then, the intra and inter VH-VL binding energies are calculated and sorted for optimized humanized antibody. The resulted humanized antibodies are ranked and compared to experimental dataset, which confirms the validity of humanized antibody design by intra and inter VH-VL binding energy.

## INTRODUCTION

Computer aided antibody humanization has become an efficient and rapid routine process[Tsuchiya and Mizuguchi, 2016; Nowak et al., 2016; Nguyen et al., 2017; Regep et al., 2017; Kuroda et al., 2020; Sawant et al., 2020], with which many humanized antibodies have been developed[Fransson et al., 2010; Apgar et al., 2016; Margreitter et al., 2016]. General process of computer aided antibody humanization includes complementarity determining region (CDR) grafting and human germline framework homology modeling[Haidar et al., 2012; Hanf et al., 2014; Choi et al., 2016; Kurella et al., 2018; Chowdhury et al., 2018; Zhang et al., 2021]. Here we report a new process of optimized antibody humanization. In this process, the CDR of murine antibody was first grafted onto homology antibody structure by loop modeling and side chain packing. Then, CDR grafted antibody structure was humanized by human germline framework. All humanized antibody structure developed in this work has been released on website [www.antibodydesign.top; www.antibodyprediction.top] and the intra and inter VH-VL binding energy were calculated and sorted to select best humanized antibody.

## METHOD

The CDR regions of user inputted non-human VH and VL sequences were automatically defined by program. Then, the proper homologous antibody structures were selected to build CDR grafted antibody by loop modeling and side chain packing. The grafted antibody structures were energy minimized and then used to build humanized antibody by selecting paired VH-VL human germline framework. The humanized antibodies were optimized by loop modeling and side chain packing. Finally, the intra and inter VH-VL binding energies of humanized antibodies were calculated and sorted to select best human germline framework(Figure 1). The humanization pipeline was provided as a webserver [www.antibodydesign.top; www.antibodyprediction.top]. The details of our pipeline are discussed in the following.

**Figure 1.**
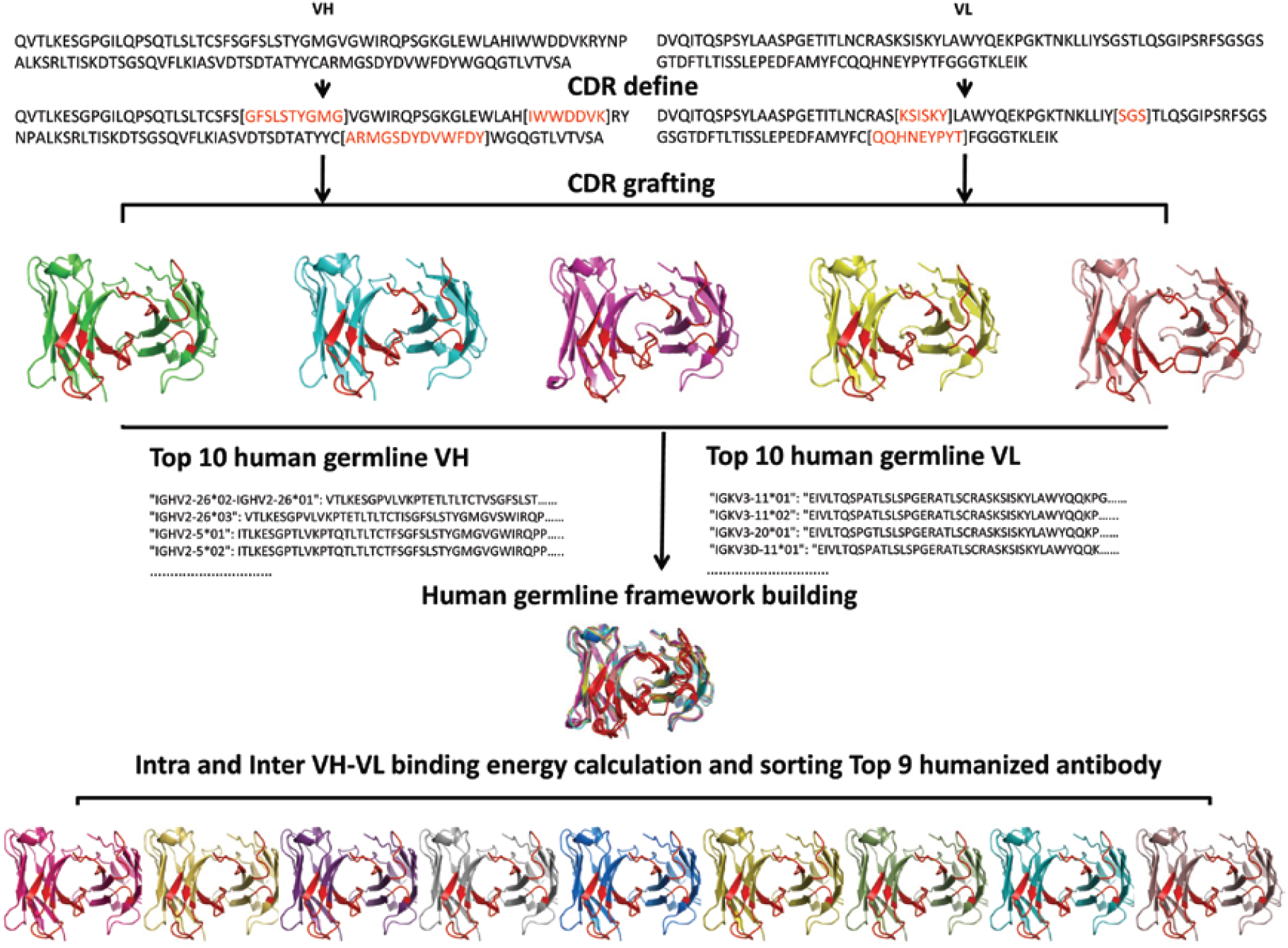
Schematics of the computer aided antibody humanization pipeline. The CDR regions of user inputted non-human VH and VL sequences were automatically defined by program. Then, the proper homologous antibody structures were selected to build CDR grafted antibody by loop modeling and side chain packing. CDR grafted antibody structure sequence was mutated to same as human germline framework sequence. Mutated residues were optimized by loop modeling and side chain packing. Then, intra and inter VH-VL binding energy was calculated. Where template intra VH and VL binding energy larger than -900kal/mol or inter VH-VL binding energy larger than -25kcal/mol was rejected. The amount of intra and inter-VH-VL binding energy was sorted to select best humanized antibody.

### Protein loop modeling

Protein backbone conformation was generated from the starting structure using the inverse kinematics method [Coutsias et al., 2004; Milgram et al., 2008; Mak et al., 2011]. These conformations could be sampled efficiently by torsion angle of the backbone from initial structure (where bond angles and lengths were fixed). For example, using a native peptide as the starting structure, the inverse kinematics methods can be depicted as:

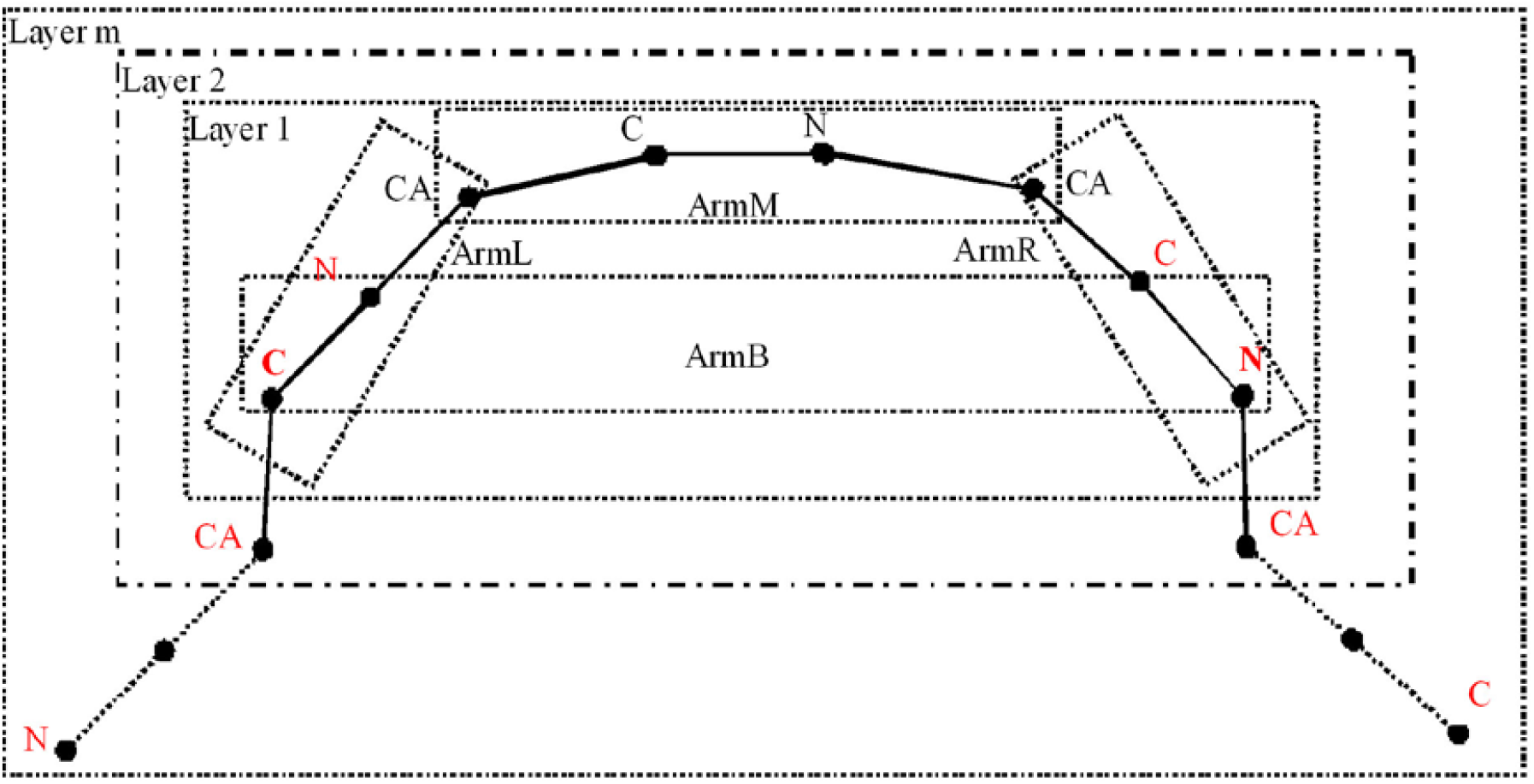
Layered inverse kinematics method

N, CA and C. denotes the corresponding backbone atoms. Layer 1 was defined as 8 atoms and marked with box by dashed line. Atoms marked in red were restrained. The restraints would extend till layer m was calculated. In layer 1, the three atoms on the left defined ArmL coordination system, and the three atoms on the right defined ArmR coordination system, the four atoms on the center defined ArmM coordination system, and the terminal atoms defined ArmB coordination system. Other layers were also defined in similar strategy.

Every layer represents a 6R inverse kinematics problem as below:

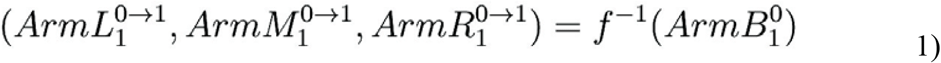

Here 0→1 denotes the conformation transition from the starting structure to the first frame. This problem has been studied extensively [Coutsias et al., 2004; Milgram et al., 2008; Mak et al., 2011]. Conformational transition of other layers could be calculated with computational scheme via

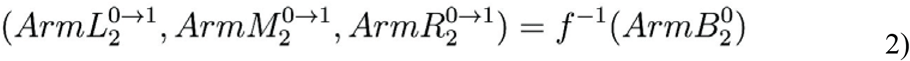

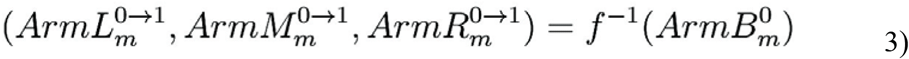

Where *m* represents layer *m*. all the conformation generated from 0→1 transition would become the starting conformation for 1→2 transition. Therefore, the *j*thframe conformation transition can be represented as:

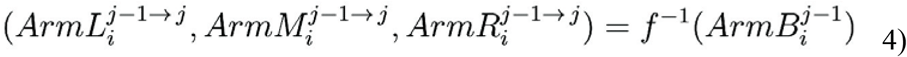

*i* represents the *i*th layer, *j* represents the *j*th frame. A backbone conformation will be rejected if the same conformation has already been generated in any previous computation steps or the conformation is physically impossible [Chen et al., 2010]. Allowable backbone conformation will be accepted for side chain packing as described below.

### Protein side chain packing

Protein side chains were constructed on backbone according to side chain torsion angle which was provided by rotamer library [Shapovalov et al., 2011]. Then, interaction energies between rotamers and template (backbone atoms only) were calculated by *van der Waals* and electrostatic interactions, rotamers with any clashes will be removed. If there were no rigid rotamers, flexible side chain will be constructed. Pairing interaction energy between rotamers at different residue position was evaluated, high energy rotamers were eliminated by dead-end elimination (DEE)method [Goldstein et al., 1994], and then global minimum energy conformation (GMEC) was determined by branch-and-terminate method [Gordon et al, 1999]. This packing method was tested on a 379 data set, the accuracy of χ_1_was 85%,χ_1_+χ_2_was 74% which was comparable with SCWRL4 [Krivov et al., 2009].

For the obtained conformations with side chain packed, those conformation will be rejected if the distance of any non-covalent bond heavy atom pair was less than 2.0Å, otherwise the binding energy will be calculated. If the calculated binding energy was larger than zero, the conformation will also be rejected. Finally, all accepted conformations were saved for later analysis.

### Antibody humanization

CDR grafted antibody structure sequence was mutated to be the same as human germline framework sequence. Mutated residues were optimized by loop modeling and side chain packing strategy described above. Then, the intra and inter VH-VL binding energies were calculated. Where template intra VH and VL binding energy larger than -900kal/mol or inter VH-VL binding energy larger than -25kcal/mol was rejected. The calculated intra and inter-VH-VL binding energies were then sorted to select best humanized antibody.

## RESULT and DISCUSSION

### Intra and Inter VH-VL binding energy of humanized antibody crystal structure

The intra and inter VH-VL binding energy of humanized antibody was calculated from the crystal structures and shown in Table 1. The intra VH and VL binding energies range from -1000kcal/mol to -1400kcal/mol, and the inter VH-VL binding energies range from -40kcal/mol to -60kcal/mol. These values are in good agreement with antibody structure database(their intra VH and VL binding energy ranged from -900kcal/mol to -1500kcal/mol, inter VH-VL binding energy ranged from - 25kc al/mol to -60kc al/mol).

**Table 1.**
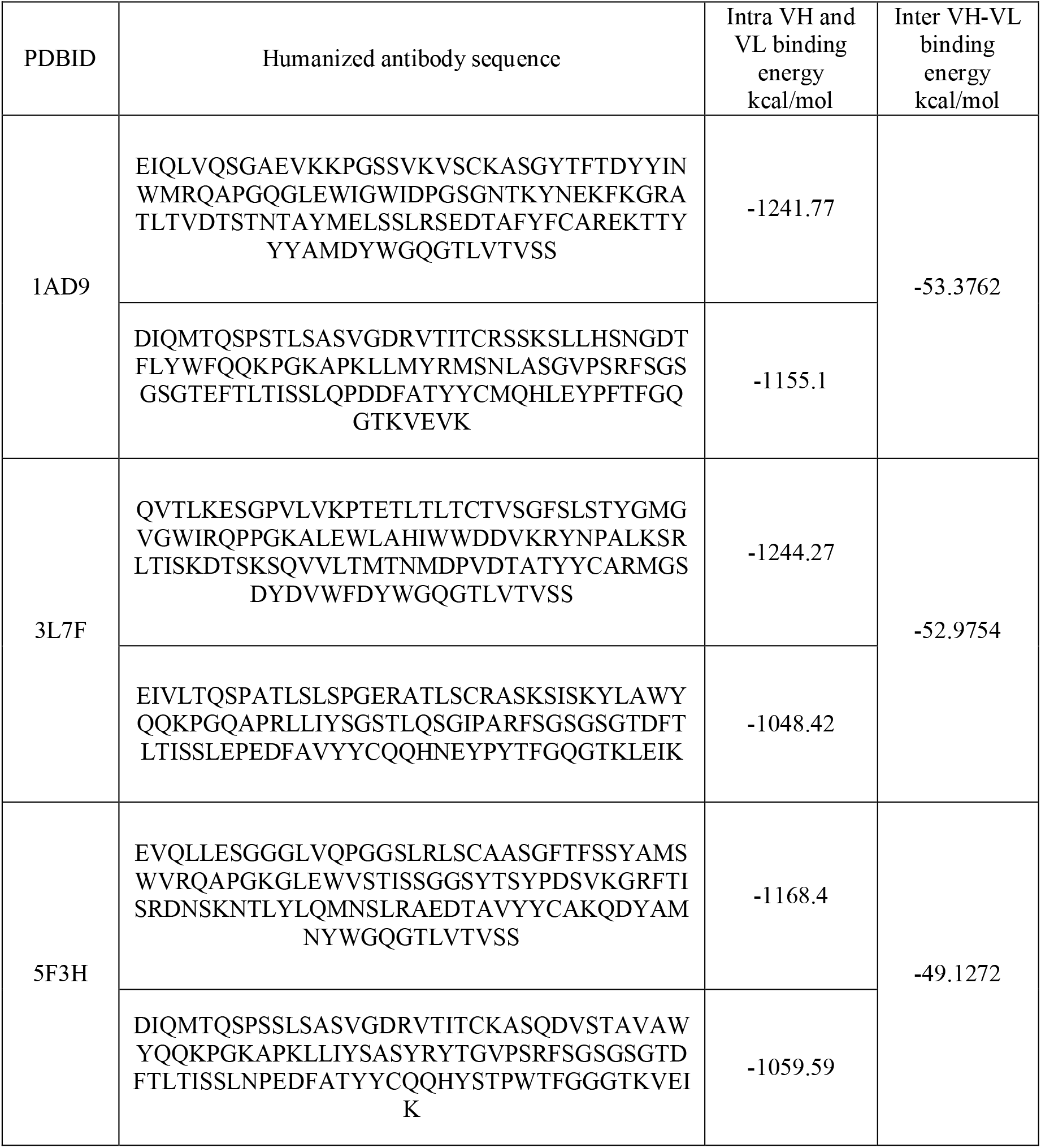
Humanized antibody crystal structure template calculated binding energy

### Murine and other antibody crystal structures as template for humanization

Murine or other antibody crystal structures were selected as template for humanization (CDR have already been in antibody crystal structure). Then, the template structures were humanized by paired VH-VL human germline framework. Humanized antibody structures were built by loop modeling and side chain packing. Then, Intra and inter VH-VL binding energies of humanized antibody were calculated and shown in Table 2. Intra VH and VL binding energies range from -900kcal/mol to -1400kcal/mol, inter VH-VL binding energies range from -40kcal/mol to -60 kcal/mol.

**Table 2.**
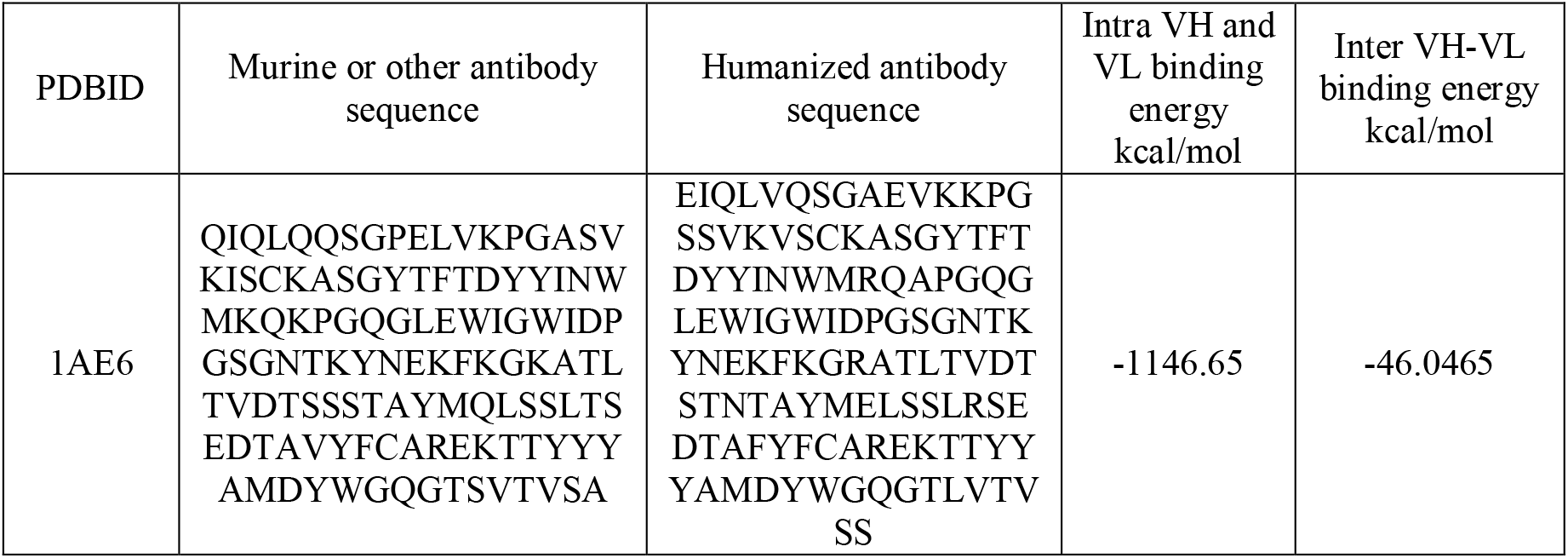
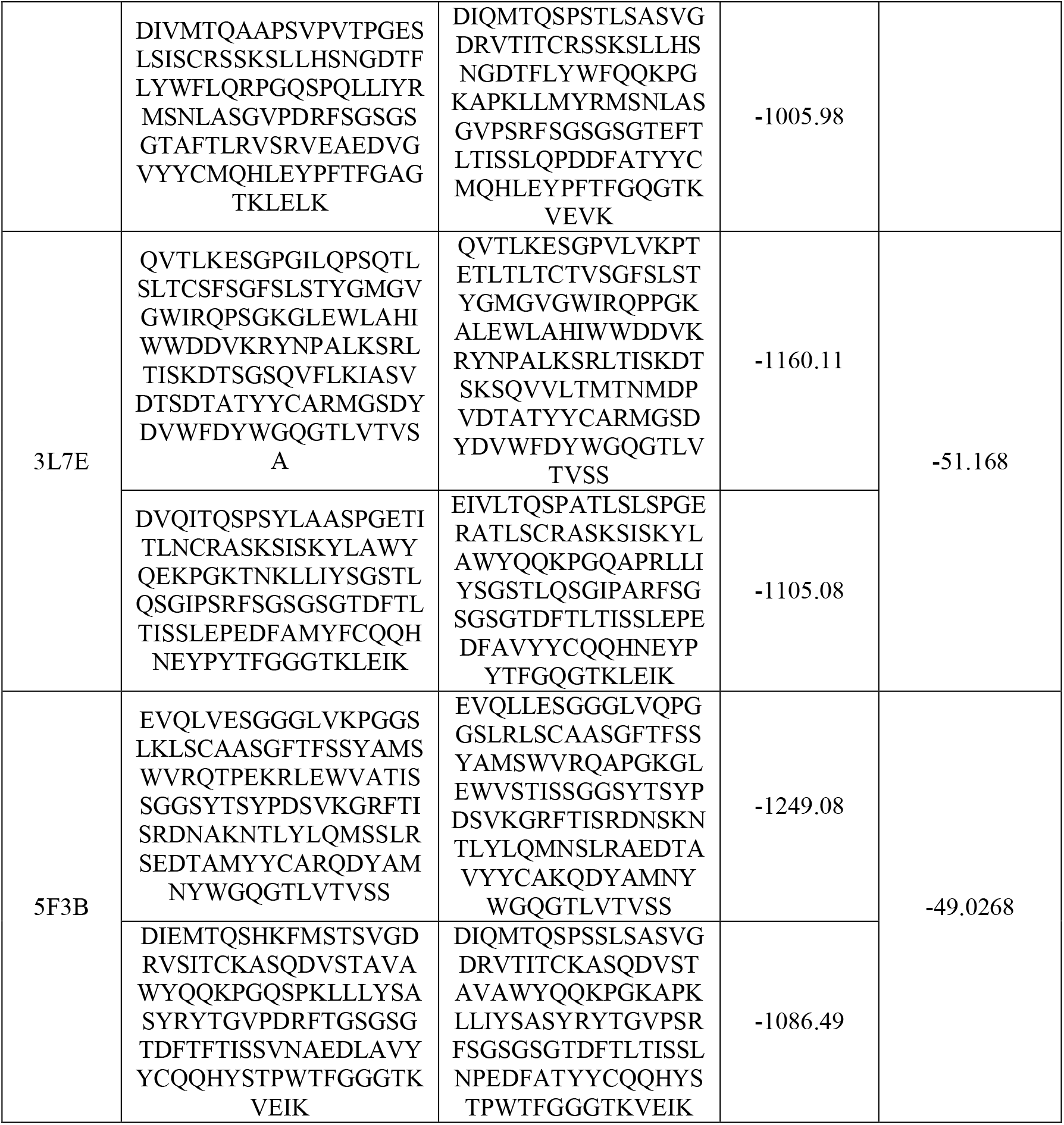
Murine or other antibody crystal structure template calculated binding energy

### Non-human derived antibody crystal structures as template for humanization

Non-human derived antibody crystal structures were selected by sequence similarity with the user inputted VH and VL sequences. Non-human antibodies were selected as templates of CDR grafting, loop modeling and side chain packing. Optimized CDR grafted antibody structures were humanized by paired VH-VL human germline framework. Humanized antibody structures were optimized by loop modeling and side chain packing. Then, the intra and inter VH-VL binding energies were calculated and shown in Table 3. The intra VH and VL binding energies range from -900kcal/mol to 1300kcal/mol, the inter VH-VL binding energies range from - 30kc al/mol to -55 kcal/mol.

**Table 3.**
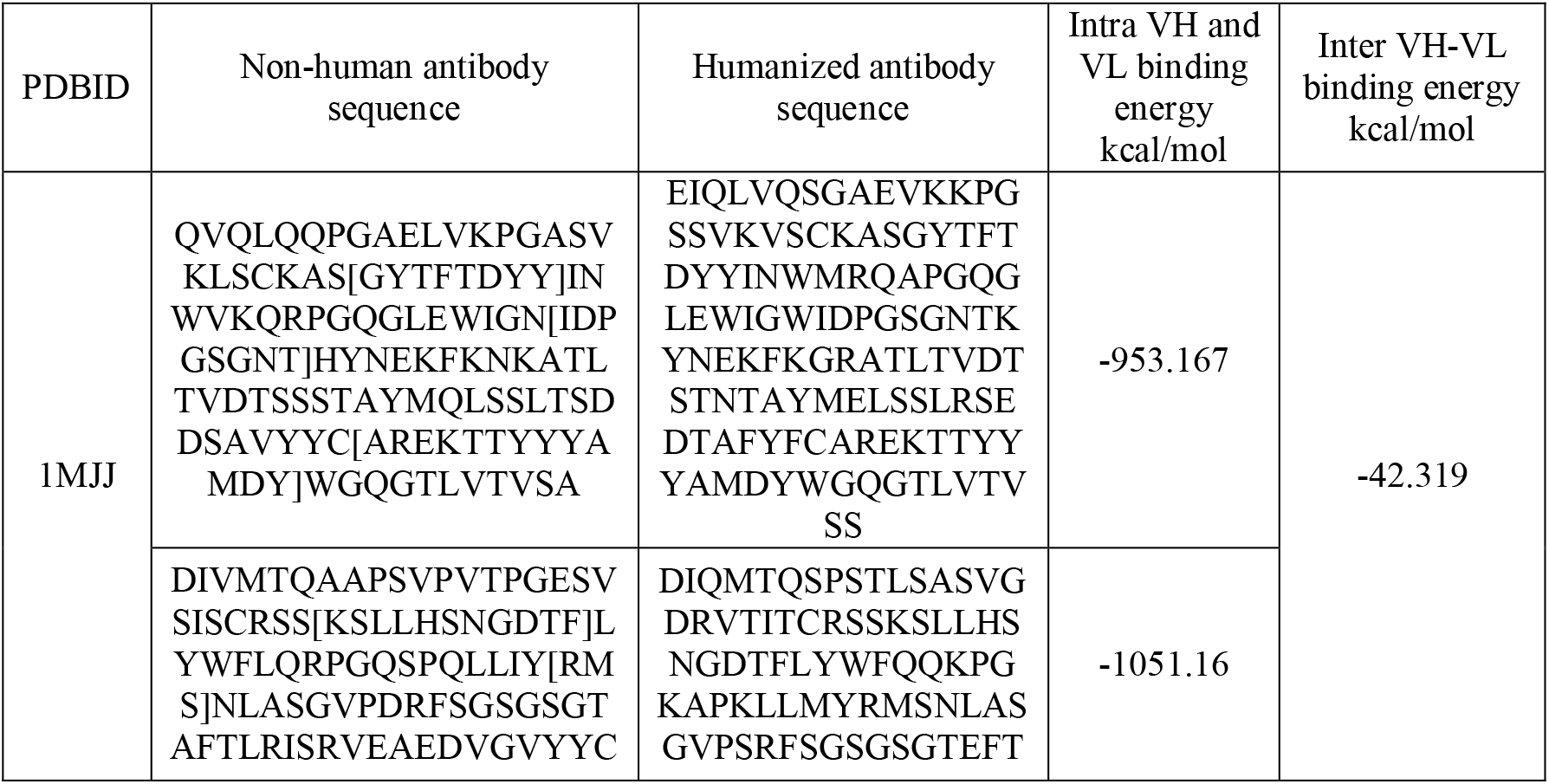
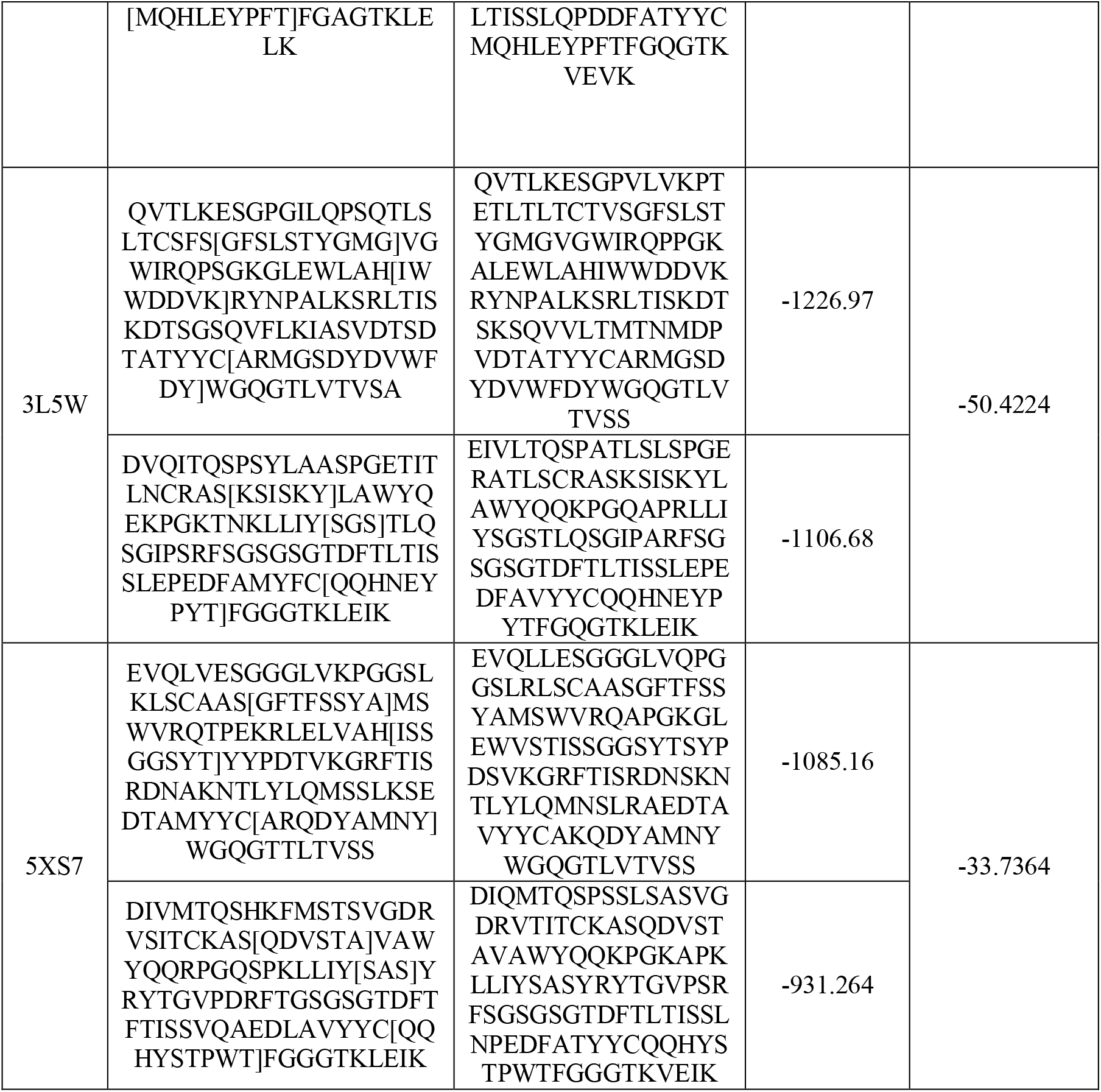
Non-human antibody crystal structure template calculated binding energy

### Humanized antibody ranking by template source from humanized antibody crystal structure

The 10 toppest VH and VL human germline framework sequences were selected by sequence similarity from the humanized antibody crystal structures. As an example, in Table 4, VH and VL human germline frameworks were combined randomly to build humanized antibody structure by template source from humanized antibody crystal structure. Humanized antibody structures were optimized and the intra and inter VH-VL binding energies were calculated. As shown in Table 5-7, rank1-9 humanized antibodies were selected by the intra and inter VH-VL binding energies.

**Table 4.** Top 10 VH and VL human germline framework

| Rank | 1AD9 |  | 3L7F |  | 5F3H |  |
| --- | --- | --- | --- | --- | --- | --- |
| 1 | 1AD9H | 1AD9L | 3L7FH | 3L7FL | 5F3HH | 5F3HL |
| 2 | IGHV1-<br>3*01 | IGKV1-<br>5*01 | IGHV2-26*02-IGHV2-26*01 | IGKV3-11*01 | IGHV3-23*03 | IGKV1D-12*02-IGKV1D-12*01-IGKV1-12*02-IGKV1-12*01 |
| 3 | IGHV1-18*04-IGHV1-18*01 | IGKV1-<br>5*03 | IGHV2-26*03 | IGKV3D-<br>7*01-IGKV3/OR2-268*02-IGKV3/OR2-268*01 | IGHV3-23*05 | IGKV1-6*02-IGKV1-6*01 |
| 4 | IGHV1-<br>69D*01-<br>IGHV1-<br>69*13-<br>IGHV1-<br>69*12- | IGKV1-<br>5*02 | IGHV2-5*02 | IGKV3D-11*03 | IGHV3-23*04 | IGKV1D-13*02-IGKV1D-13*01-IGKV1-13*02 |
|  | IGHV1-69*01 |  |  |  |  |  |
| 5 | IGHV1-69*05 | IGKV1-17*01 | IGHV2-5*01 | IGKV3D-11*01 | IGHV3-23D*01-IGHV3-23*01 | IGKV1D-39*01-IGKV1-39*01 |
| 6 | IGHV1-69*16 | IGKV1-16*01 | IGHV2-5*09 | IGKV3D-11*02 | IGHV3-53*02 | IGKV1-39*02 |
| 7 | IGHV1-69*15 | IGKV1-NL1*01 | IGHV2-5*06-IGHV2-5*05 | IGKV3-11*02 | IGHV3-23*02 | IGKV1-NL1*01 |
| 8 | IGHV1-69*11 | IGKV1-17*03 | IGHV2-5*04 | IGKV3D-20*02 | IGHV3-66*04-IGHV3-66*02-IGHV3-66*01 | IGKV1-5*01 |
| 9 | IGHV1-69*17 | IGKV1-9*01 | IGHV2-5*08 | IGKV3-20*01 | IGHV3-66*03 | IGKV1-5*03 |
| 10 | IGHV1-69*14-IGHV1-69*06 | IGKV1-17*02 | IGHV2/OR16-5*01 | IGKV3D-15*01-IGKV3-15*01 | IGHV3-53*01 | IGKV1-27*01 |

**Table 5.** Top 9 humanized antibody template with 1AD9

| Rank | HL Gene | H intra kcal/mol | L intra kcal/mol | HL inter kcal/mol |
| --- | --- | --- | --- | --- |
| 1 | IGHV1-3*01-1AD9L | -1264.06 | -1174.53 | -52.1395 |
| 2 | IGHV1-3*01-IGKV1-NL1*01 | -1273.36 | -1133.73 | -51.4032 |
| 3 | IGHV1-69*11-1AD9L | -1234.91 | -1170.49 | -50.8565 |
| 4 | IGHV1-69*15-1AD9L | -1234.91 | -1170.49 | -50.8565 |
| 5 | 1AD9H-1AD9L | -1241.77 | -1155.1 | -53.3762 |
| 6 | IGHV1-69*05-1AD9L | -1218.29 | -1171.59 | -50.3606 |
| 7 | IGHV1-69*16-1AD9L | -1218.29 | -1171.59 | -50.3606 |
| 8 | 1AD9H-IGKV1-NL1*01 | -1249.31 | -1135.32 | -52.2981 |
| 9 | IGHV1-69D*01-IGHV1-69*13-IGHV1-69*12-IGHV1-69*01-1AD9L | -1215.11 | -1171.07 | -50.4236 |

**Table 6.** Top 9 humanized antibody template with 3L7F

| Rank | HL Gene | H intra kcal/mol | L intra kcal/mol | HL inter kcal/mol |
| --- | --- | --- | --- | --- |
| 1 | 3L7FH-IGKV3D-20*02 | -1245.37 | -1068.45 | -52.9509 |
| 2 | 3L7FH-IGKV3-11*02 | -1246.3 | -1064.82 | -52.8501 |
| 3 | 3L7FH-IGKV3-11*01 | -1246.2 | -1065.4 | -49.4023 |
| 4 | 3L7FH-IGKV3D-15*01-IGKV3-15*01 | -1245.52 | -1061.7 | -49.9812 |
| 5 | 3L7FH-IGKV3-20*01 | -1246.05 | -1054.99 | -52.8499 |
| 6 | 3L7FH-3L7FL | -1244.27 | -1048.42 | -52.9754 |
| 7 | 3L7FH-IGKV3D-7*01-IGKV3/OR2-268*02-IGKV3/OR2-268*01 | -1245.91 | -1040.16 | -50.3278 |
| 8 | IGHV2-5*09-IGKV3D-20*02 | -1068.69 | -1075.64 | -51.972 |
| 9 | IGHV2-5*09-IGKV3-11*02 | -1068.45 | -1074.8 | -52.1395 |

**Table 7.**
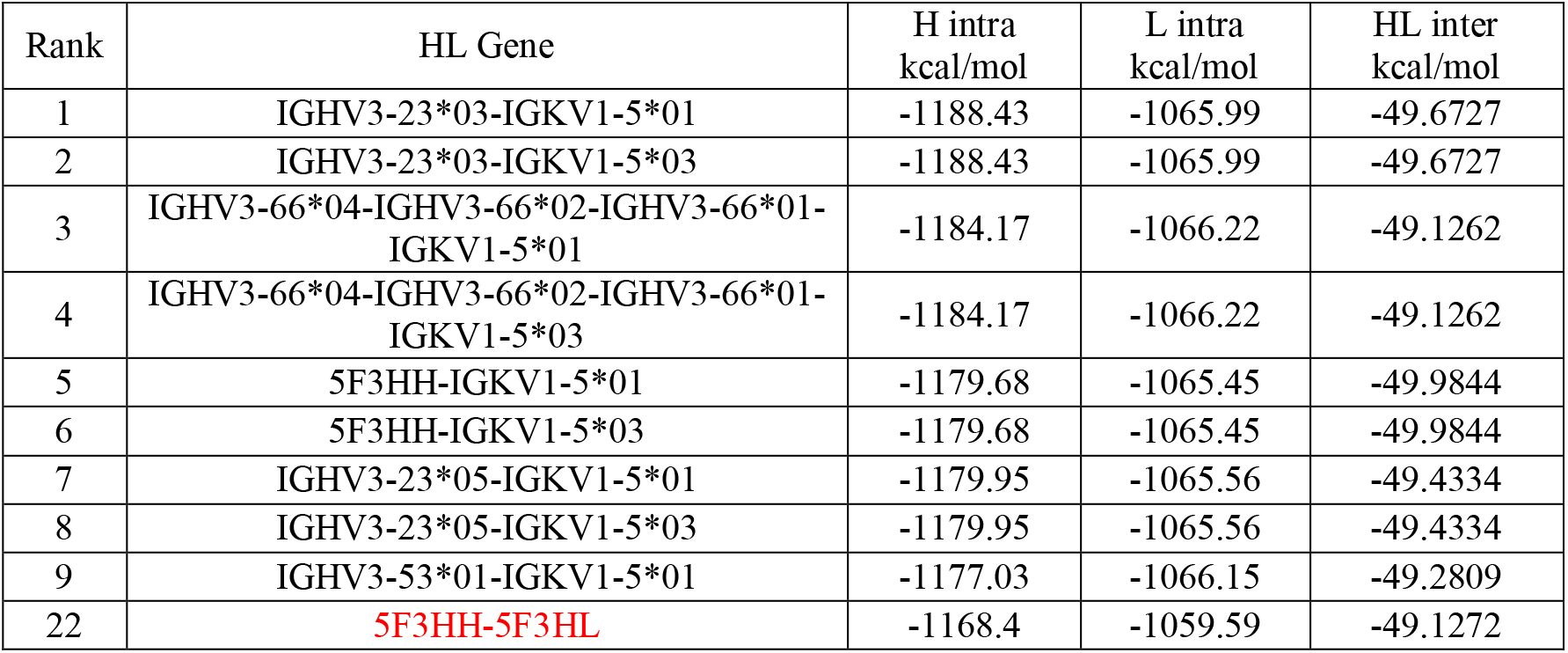
Top 9 humanized antibody template with 5F3H

### Humanized antibody ranking by template source from murine and other antibody crystal structure

Murine and other source antibody crystal structures were selected as templates to build humanized antibodies (CDR has already been in antibody crystal structure). Then human germline frameworks in Table 4 were randomly combined for humanization. Humanized antibodies were optimized by loop modeling and side chain packing. Then, the intra and inter VH-VL binding energies of the optimized humanized antibodies were calculated for sorting of the top 9 humanized antibodies. Some of the humanized antibodies VH and VL framework (shown in Table 8-10) could also be found in Table 5-7. Therefore, the intra and inter VH-VL binding energies could be used for the selection of the best humanized antibody source from murine and other antibody crystal structures.

**Table 8.** Top 9 humanized antibody template with 1AE6

| Rank | HL Gene | H intra kcal/mol | L intra kcal/mol | HL inter kcal/mol |
| --- | --- | --- | --- | --- |
| 1 | IGHV1-3*01-1AD9L | -1127.61 | -1047.12 | -53.0664 |
| 2 | 1AD9H-IGKV1-5*01 | -1143.12 | -1032.38 | -48.7873 |
| 3 | 1AD9H-IGKV1-5*03 | -1143.12 | -1032.38 | -48.7873 |
| 4 | 1AD9H-IGKV1-5*02 | -1140.71 | -1029.61 | -50.2803 |
| 5 | IGHV1-3*01-IGKV1-5*01 | -1125.35 | -1033.61 | -52.8156 |
| 6 | IGHV1-3*01-IGKV1-5*03 | -1125.35 | -1033.61 | -52.8156 |
| 7 | 1AD9H-IGKV1-17*02 | -1143.97 | -1026.23 | -46.9193 |
| 8 | 1AD9H-IGKV1-17*01 | -1147.72 | -1022.63 | -46.7637 |
| 9 | IGHV1-3*01-IGKV1-5*02 | -1128.03 | -1031.66 | -52.033 |

**Table 9.** Top 9 humanized antibody template with 3L7E

| Rank | HL Gene | H intra kcal/mol | L intra kcal/mol | HL inter kcal/mol |
| --- | --- | --- | --- | --- |
| 1 | IGHV2-5*08-IGKV3D-20*02 | -1185.35 | -1115.58 | -47.9664 |
| 2 | IGHV2-5*08-3L7FL | -1182.2 | -1108.66 | -50.4048 |
| 3 | IGHV2-5*08-IGKV3-11*02 | -1185.55 | -1107.84 | -48.2813 |
| 4 | IGHV2-5*08-IGKV3-20*01 | -1185.31 | -1107.56 | -48.0529 |
| 5 | IGHV2-5*04-IGKV3D-20*02 | -1175.21 | -1114.91 | -48.0891 |
| 6 | IGHV2-5*01-IGKV3D-20*02 | -1172.76 | -1115.2 | -48.1578 |
| 7 | IGHV2-5*02-IGKV3D-20*02 | -1172.76 | -1115.2 | -48.1578 |
| 8 | IGHV2-5*08-IGKV3-11*01 | -1185.57 | -1102.47 | -48.0565 |
| 9 | IGHV2-5*04-IGKV3-11*02 | -1176.39 | -1109.7 | -48.1239 |

**Table 10.** Top 9 humanized antibody template with 5F3B

| Rank | HL Gene | H intra kcal/mol | L intra kcal/mol | HL inter kcal/mol |
| --- | --- | --- | --- | --- |
| 1 | IGHV3-53*01-5F3HL | -1270.01 | -1086.12 | -47.9012 |
| 2 | IGHV3-66*03-5F3HL | -1270.01 | -1086.12 | -47.9012 |
| 3 | IGHV3-66*04-IGHV3-66*02-IGHV3-66*01-5F3HL | -1268.52 | -1085.73 | -48.038 |
| 4 | IGHV3-23*03-5F3HL | -1266.15 | -1086.16 | -48.9234 |
| 5 | IGHV3-53*02-5F3HL | -1262.08 | -1086.12 | -48.8344 |
| 6 | IGHV3-23*03-IGKV1-5*01 | -1265.61 | -1084.3 | -46.7451 |
| 7 | IGHV3-23*03-IGKV1-5*03 | -1265.61 | -1084.3 | -46.7451 |
| 8 | IGHV3-66*04-IGHV3-66*02-IGHV3- | -1267.16 | -1084.06 | -45.8075 |
|  | 66*01-IGKV1-5*01 |  |  |  |
| 9 | IGHV3-66*04-IGHV3-66*02-IGHV3-66*01-IGKV1-5*03 | -1267.16 | -1084.06 | -45.8075 |

### Humanized antibody ranking by template source from non-human antibody crystal structure

Non-human derived antibody structures were selected by sequence similarity with the user inputted VH and VL sequences. Then, CDR grafted antibodies were built by CDR grafting, loop modeling and side chain packing. The optimized CDR grafted antibodies were humanized by human germline framework shown in Table 4. Humanized antibodies were optimized by loop modeling and side chain packing. Then, the intra and inter VH-VL binding energies of the humanized antibodies were calculated for sorting the top 9 humanized antibody. As shown in Table 11-13, some of the rank 1-9 humanized antibody VH-VL genes could also be found in Table 5-7. Therefore, the intra and inter VH-VL binding energies could also be used for selection of the best humanized antibody source from non-human crystal structure.

**Table 11.** Top 9 humanized antibody template with 1MJJ

| Rank | HL Gene | H intra kcal/mol | L intra kcal/mol | HL inter kcal/mol |
| --- | --- | --- | --- | --- |
| 1 | IGHV1-3*01-1AD9L | -957.923 | -1049.05 | -47.2008 |
| 2 | IGHV1-3*01-IGKV1-5*02 | -953.934 | -1043.96 | -50.0913 |
| 3 | IGHV1-3*01-IGKV1-16*01 | -954.108 | -1044.86 | -49.342 |
| 4 | IGHV1-3*01-IGKV1-9*01 | -950.754 | -1046.37 | -48.7738 |
| 5 | IGHV1-3*01-IGKV1-5*01 | -953.907 | -1040.09 | -49.9078 |
| 6 | IGHV1-3*01-IGKV1-5*03 | -953.907 | -1040.09 | -49.9078 |
| 7 | 1AD9H-1AD9L | -953.167 | -1051.16 | -42.319 |
| 8 | 1AD9H-IGKV1-9*01 | -951.375 | -1050.56 | -41.6717 |
| 9 | 1AD9H-IGKV1-5*01 | -950.974 | -1041.92 | -46.0397 |

**Table 12.** Top 9 humanized antibody template with 3L5W

| Rank | HL Gene | H intra kcal/mol | L intra kcal/mol | HL inter kcal/mol |
| --- | --- | --- | --- | --- |
| 1 | 3L7FH-IGKV3-11*02 | -1220.15 | -1120.73 | -54.1098 |
| 2 | 3L7FH-IGKV3-11*01 | -1218.71 | -1110.03 | -55.5043 |
| 3 | 3L7FH-IGKV3D-20*02 | -1224.81 | -1113.4 | -49.979 |
| 4 | 3L7FH-3L7FL | -1226.97 | -1106.68 | -50.4224 |
| 5 | 3L7FH-IGKV3-20*01 | -1224.62 | -1102.52 | -52.3325 |
| 6 | IGHV2-5*09-IGKV3-11*02 | -1198.87 | -1122.39 | -53.9434 |
| 7 | IGHV2-5*01-IGKV3-11*02 | -1197.92 | -1122.18 | -53.0765 |
| 8 | IGHV2-5*02-IGKV3-11*02 | -1197.92 | -1122.18 | -53.0765 |
| 9 | IGHV2-5*04-IGKV3-11*02 | -1194.83 | -1122.41 | -53.9548 |

**Table 13.** Top 9 humanized antibody template with 5XS7

| Rank | HL Gene | H intra kcal/mol | L intra kcal/mol | HL inter kcal/mol |
| --- | --- | --- | --- | --- |
| 1 | IGHV3-23*04-5F3HL | -1104.9 | -929.997 | -36.4064 |
| 2 | IGHV3-23D*01-IGHV3-23*01-5F3HL | -1103.05 | -931.231 | -36.5693 |
| 3 | IGHV3-23*05-5F3HL | -1101.42 | -932.443 | -36.3889 |
| 4 | IGHV3-23*02-5F3HL | -1100.79 | -929.255 | -36.3646 |
| 5 | IGHV3-53*01-5F3HL | -1097.44 | -931.202 | -36.9109 |
| 6 | IGHV3-66*03-5F3HL | -1097.44 | -931.202 | -36.9109 |
| 7 | IGHV3-66*04-IGHV3-66*02-IGHV3-66*01-5F3HL | -1095.53 | -929.957 | -37.0653 |
| 8 | IGHV3-53*02-5F3HL | -1095.15 | -930.529 | -36.868 |
| 9 | IGHV3-23*03-5F3HL | -1093.12 | -932.089 | -36.8587 |

## CONCLUSION

The user inputted VH-VL sequences were used to search suitable antibody crystal structures. These antibody structures can be used as templates for CDR grafting, loop modeling and side chain packing. Optimized CDR grafted antibodies were humanized by human germline framework. Humanized antibodies were then optimized by loop modeling and side chain packing. Finally, the intra and inter VH-VL binding energies were calculated and sorted for the selection of the best humanized antibody. Our results indicate that the intra and inter VH-VL binding energies could be used to rank humanized antibody.

## ACKNOWLEDGMENTS

This work was supported by National High-tech R&D Program of China (863 Program) 2014AA021103. The author would like to thank Dr. Jizhong Lou for helpful discussions and revising the article.

## REFERENCE

1. Apgar JR, Mader M, Agostinelli R, Benard S, Bialek P, Johnson M, Gao Y, Krebs M, Owens J, Parris K, St Andre M, Svenson K, Morris C, Tchistiakova L. (2016) Beyond CDR-grafting: Structure-guided humanization of framework and CDR regions of an anti-myostatin antibody. MAbs. 8(7):1302–1318..

2. Choi Y, Ndong C, Griswold KE, Bailey-Kellogg C. (2016) Computationally driven antibody engineering enables simultaneous humanization and thermostabilization. Protein Eng Des Sel., 29(10):419–426.

3. Chowdhury R, Allan MF, Maranas CD. (2018) OptMAVEn-2.0: De novo Design of Variable Antibody Regions against Targeted Antigen Epitopes. Antibodies (Basel)., 7(3):23.

4. Fransson J, Teplyakov A, Raghunathan G, Chi E, Cordier W, Dinh T, Feng Y, Giles-Komar J, Gilliland G, Lollo B, Malia TJ, Nishioka W, Obmolova G, Zhao S, Zhao Y, Swanson RV, Almagro JC. (2010) Human framework adaptation of a mouse anti-human IL-13 antibody. J Mol Biol.,398(2):214–31.

5. Haidar JN, Yuan QA, Zeng L, Snavely M, Luna X, Zhang H, Zhu W, Ludwig DL, Zhu Z. (2012) A universal combinatorial design of antibody framework to graft distinct CDR sequences: a bioinformatics approach. Proteins.,80(3):896–912.

6. Hanf KJ, Arndt JW, Chen LL, Jarpe M, Boriack-Sjodin PA, Li Y, van Vlijmen HW, Pepinsky RB, Simon KJ, Lugovskoy A. (2014) Antibody humanization by redesign of complementarity-determining region residues proximate to the acceptor framework. Methods., 65(1):68–76.

7. Kuroda D, Tsumoto K.(2020) Engineering Stability, Viscosity, and Immunogenicity of Antibodies by Computational Design. J Pharm Sci., 109(5):1631–1651.

8. Kurella VB, Gali R.(2018) Antibody Design and Humanization via In Silico Modeling. Methods Mol Biol., 1827:3–14.

9. Margreitter C, Mayrhofer P, Kunert R, Oostenbrink C.(2016) Antibody humanization by molecular dynamics simulations-in-silico guided selection of critical backmutations. J Mol Recognit,29(6):266–75.

10. Nguyen MN, Pradhan MR, Verma C, Zhong P.(2017) The interfacial character of antibody paratopes: analysis of antibody-antigen structures. Bioinformatics., 33(19):2971–2976.

11. Nowak J, Baker T, Georges G, Kelm S, Klostermann S, Shi J, Sridharan S, Deane CM. (2016) Length-independent structural similarities enrich the antibody CDR canonical class model. MAbs.,8(4):751–60.

12. Regep C, Georges G, Shi J, Popovic B, Deane CM. (2017) The H3 loop of antibodies shows unique structural characteristics. Proteins., 85(7):1311– 1318.

13. Sawant MS, Streu CN, Wu L, Tessier PM. (2020) Toward Drug-Like Multispecific Antibodies by Design. Int J Mol Sci., 21(20):7496.

14. Tsuchiya Y, Mizuguchi K. (2016) The diversity of H3 loops determines the antigen-binding tendencies of antibody CDR loops. Protein Sci., 25(4):815– 25.

15. Zhang CH, Kim K, Jin Z, Zheng F, Zhan CG. (2021) Systematic Structure-Based Virtual Screening Approach to Antibody Selection and Design of a Humanized Antibody against Multiple Addictive Opioids without Affecting Treatment Agents Naloxone and Naltrexone. ACS Chem Neurosci., 12(1):184–194.

16. Shapovalov, M., & Dunbrack, R. (2011) A smoothed backbone-dependent rotamer library for proteins derived from adaptive kernel density estimates and regressions. Structure., 19(6), 844–858.

17. Goldstein RF. (1994) Efficient Rotamer Elimination Applied to Protein Side-Chains and Related Spin Glasses. Biophysical Journal., 66:1335–1340.

18. Gordon DB., & Mayo SL. (1999) Branch-and-terminate: a combinatorial optimization algorithm for protein design. Structure., 7(9), 1089–1098.

19. Krivov GG., Shapovalov MV., & Dunbrack RL. Improved prediction of protein side-chain conformations with scwrl4. Proteins Structure Function & Bioinformatics. 2009, 77(4),: 778–795.

20. Coutsias EA., Seok C., Jacobson MP., & Dill KA. (2004) A kinematic view of loop closure.. Journal of Computational Chemistry., 25(4), 510–528.

21. Milgram RJ., Liu G., & Latombe JC. (2008) On the structure of the inverse kinematics map of a fragment of protein backbone. Journal of Computational Chemistry., 29(1), 50–68.

22. Mak CH. (2011) Loops mc: an all-atom monte carlo simulation program for rnas based on inverse kinematic loop closure. Molecular Simulation., 37(7), 537–556.

23. Chen VB., Arendall WB., Headd JJ., Keedy DA., Immormino RM., & Kapral GJ., et al. (2010) Molprobity: all-atom structure validation for macromolecular crystallography. Acta Crystallogr D Biol Crystallogr., 66(1), 12–21.

